# The spatio-temporal architecture of everyday manual behavior

**DOI:** 10.1101/2022.09.21.508833

**Authors:** Daniele Sili, Chiara De Giorgi, Alessandra Pizzuti, Matteo Spezialetti, Francesco de Pasquale, Viviana Betti

**Author notes:** now, at a different institution (Chiara De Giorgi, Alessandra Pizzuti). now, at Department of Information Engineering, Computer Science and Mathematics, University of L’Aquila (Matteo Spezialetti).

## Abstract

In everyday activities, humans use a finite number of postural hand configurations, but how do they flow into each other to create sophisticated manual behavior? We hypothesized that hand movement emerges through the temporal dynamics of a set of recurrent hand shapes characterized by specific transitions. Through a sensorized glove, we collected kinematics data from thirty-six participants preparing and having breakfast in naturalistic conditions. By means of a combined PCA/clustering-based approach, we identified a repertoire of hand states and their transitions over time. We found that manual behavior can be described in space through a complex organization of basic configurations. These, even in an unconstrained experiment, recurred across subjects. A specific temporal structure, highly consistent within the sample, seems to integrate such identified hand shapes to realize skilled movements. Our findings suggest that the simplification of the motor commands unravels in the temporal dimension more than in the spatial one.

## Introduction

Motor behavior is dynamic by nature, structured in successions of simple motor acts. From learning how to use chopsticks in a Japanese restaurant, to grasping everyday objects, hand movements are organized into a precise temporal ordering. During the movement, the proper sequence of hand shapes is fundamental to the success of manipulating actions directed at daily life objects.

Postural configurations arise from both biomechanics and neural constraints defining the degrees of freedom for volitional control, hence the various ways to perform a movement ^1^. How the central nervous system controls the hand movement is still a debated issue^2, 3^, especially considering its temporal dynamics. One leading hypothesis suggests that the brain is concerned more with postures than movements^4^. A preferential encoding of time-varying postures has been recently shown in neurons of the sensorimotor cortex^5^. This is in line with classical studies showing activation of neurons in the primate frontal cortex for specific goal-oriented motor acts rather than isolated finger movements ^6, 7^.

Despite its complexity, the prevalent idea is that the human hand uses a small set of hand configurations, i.e., distinct functional modules or postural synergies, to solve different motor tasks ^8–10^. A synergy can be broadly defined as a set of muscle activations or finger joint angles recruited collectively rather than individually, for a specific motor goal^10^. They are also referred to as covariation patterns and are typically extracted from electromyography (EMG) or kinematic signals^11–13^ through Principal Component Analysis (PCA)^14^ or non-negative factorization (NMF)^15^. Using PCA, the basic assumption is that each principal component (PC), explaining the largest variance, represents a hand synergy involved with the movement. Thus far, most of the existing studies on hand synergies analyzed limited sets of static hand postures associated with stereotyped manual behaviors^13^, imagined familiar objects^10^, grasping of recalled and virtual objects^16^, or considering a predefined phase of hand movements (i.e., before contacting the object^17^ or at the end of the transport phase^10^). These studies observed that hand movements are limited to a few covariance patterns involving muscles/joints and fingers. Typically, from three to six principal components account for more than 80% of the variability of hand movements, see for example^10^, but also^18^ for a high-dimensional manifold of hand kinematics.

However, natural hand movement unfolds over time ^7, 19^. Hence, models considering both spatial and temporal architecture of the synergy are the most suitable to describe multiple muscles activation^20, 21^ or joints posture^16, 22–25^. Furthermore, everyday manual behavior involves multi-joint movements and consists of a large variety of unconstrained postures. Nevertheless, existing studies on hand dynamics mainly concern hand shaping within experimentally controlled movements^16, 22, 26^. These studies adopted a reduced dimensional space (see, for example^16, 22^) or a weighted variation of a mean (static) posture^26^. Often, they also used a small sample of participants (from four to eight, e.g.,^10, 13, 16, 17, 22, 26–28)^.

By contrast, here, we investigated how the hand movement develops when thirty-six participants spontaneously interact with various daily life objects. Natural interaction with common objects is an unconstrained paradigm, which has been previously adopted to study the statistics of natural hand movements ^27^. Specifically, we tested whether a precise temporal ordering orchestrates the dynamics of a discrete set of “hand states” during the natural interaction with common objects. The term “hand state” refers here to a hand configuration that can span a few seconds, recurs over time, and is consistently observed across participants and objects. In other words, we aim to uncover how the hand switches between states and the occurrence of these changes. Understanding how natural hand behavior builds across space and time is fundamental. It allows to unravel the link with the underlying neural dynamics^29^, and with networks allowing flexible switching within a repertoire of hand movements^30^ . Based on the idea that natural behavior can be modeled as discrete states alternating over time^31, 32^, we developed a novel approach. We modeled, natural hand movements as a set of postural states, for which we infer their occurrence and dynamic switch. Results suggest that a precise spatiotemporal architecture orchestrates natural hand behavior. However, differently from previous studies suggesting synergies as a simplified description of hand action, here we observed a complex spatial structure. By contrast, the observed temporal structure seemed to represent a simplification mechanism to orchestrate such complexity.

## Results

This study aims at identifying the most recurrent hand configurations and how they flow into each other in a large cohort of participants performing movements in a naturalistic context. When exploring their dynamics, we will refer to them as hand states. Of note, based on previous frameworks^1, 9, 10^, a synergy represents here a basic form of movement, the building-block of each postural configuration. This is defined at a spatial level as joint co-variation. The hand state refers to the composition of similar hand configurations observed among participants. It is defined based on its temporal dynamics, in terms of recurrence and switch to form the complexity of natural movement.

Thirty-six participants performed reach-to-grasp movements, entailing contact with everyday objects (Table S1), aimed at preparing and having breakfast (Fig. 1a).

**Figure 1.**
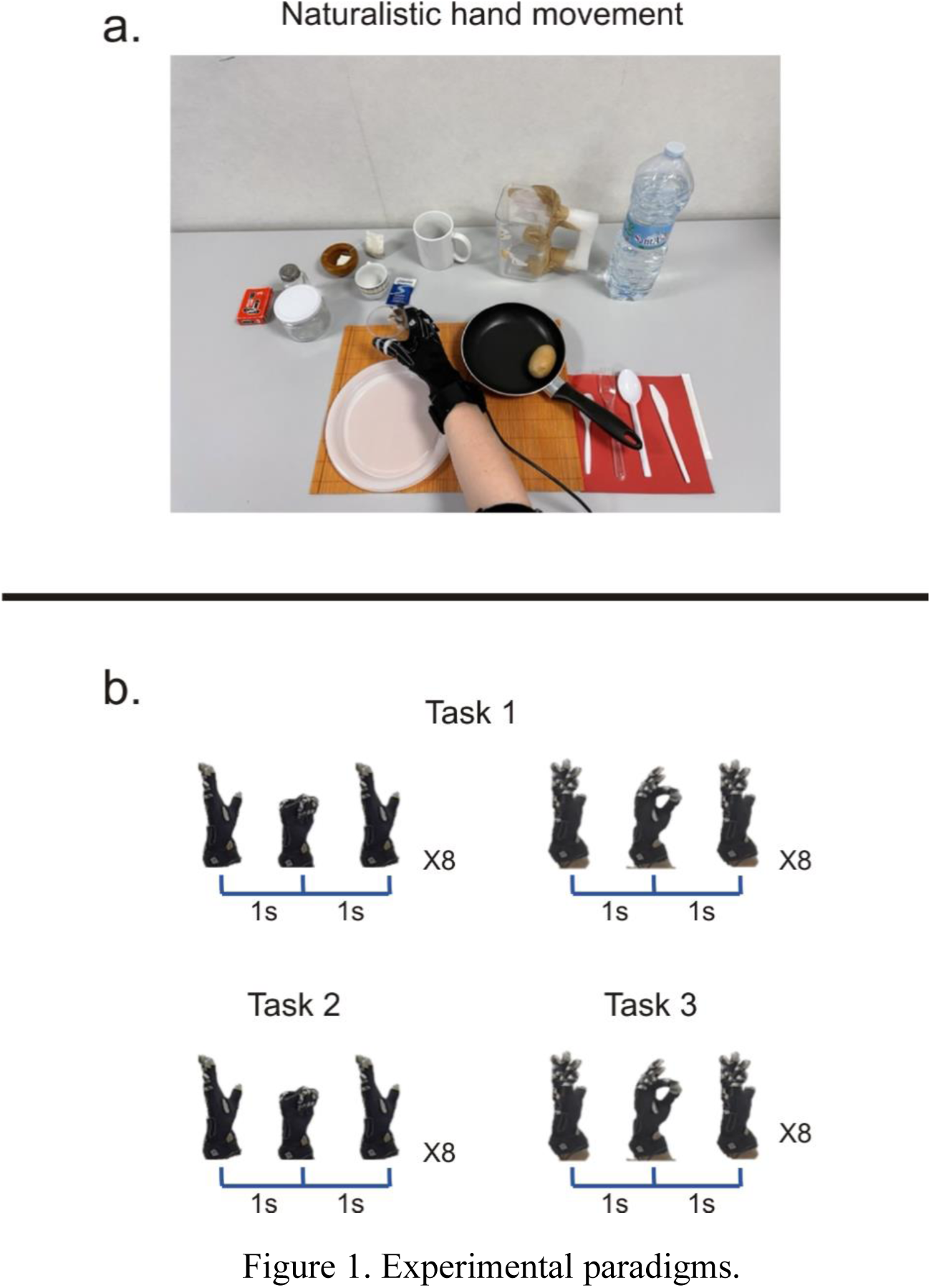
Experimental paradigms. (a) The naturalistic experiment. Participants were asked to interact with objects necessary for preparing and eating breakfast, trying to reproduce the actions they usually perform in the kitchen. Participants were seated and asked to set the table, mimic the act of cooking and preparing breakfast, eat the meal and finally clean and arrange the table to its initial state. (b) Canonical experiment. A single participant was asked to execute canonical movements (not entailing contact with the objects). Four different stereotyped movements were performed: a sequence of hand opening-closure followed by a pinch movement (task1), or hand opening-closure movement (task2) and pinch movement (task 3) in isolation. The movements were repeated 8 times for 2 seconds.

The kinematics data were acquired through a CyberGlove (see Materials and Methods). First, we identified the set of hand shapes underlying natural movements. To this aim, after data calibration and preprocessing, we performed PCA to extract postural synergies ^11, 14, 33^, separately for time windows (see Materials and Methods). In this way, we restricted hand configurations to those explaining most of the variance in the data. In Fig. S1 we report the analysis pipeline. We note that the hypothesis of a low dimensionality of the kinematics holds. As an example, we show in Fig. S1b data from a representative subject, where in the reported time window, 90% of the variance was explained by 4 PCs. The group analysis confirmed that the dimensionality of the hand configurations across all subjects and windows was smaller than the degrees of freedom (see Figure S1c, left panel and Material and Methods).

Next, we explored how the low dimensional structure of hand kinematics varies over time. This is of interest, as we considered the hand movements in all its phases, e.g., pre-shaping, transport, object contact, and action-oriented manipulation. One might expect that, although at the subject level PCs were reproducible^26^, they could serve different movements. This is in keeping with the idea that the same synergy can be shared across several distinct hand actions ^24, 34^. To address this aspect, we firstly ranked the obtained PCs in four groups based on the amount of variance explained (see Materials and Methods). Then, we clustered the obtained synergies across participants to identify the final hand states and their temporal dynamics.

### Synergies and hand shapes during canonical movements

Before applying our approach to the naturalistic paradigm, we investigated the relationship between extracted synergies and hand shapes in a controlled experiment. This was designed with canonical movements. In this way, we tested our analysis pipeline to be applied to the data collected in the naturalistic scenario (next section).

Here, a single participant performed stereotyped hand movements (not entailing contact with objects) either in sequence, i.e., hand opening-closure followed by a pinch movement (task 1), or in isolation, i.e., opening-closure (task 2), pinch movement (task 3) (Fig. 1b). In both tasks, the sequence was repeated eight times for 2 seconds (Fig. 1b). As can be noted in Figure 2a, in both tasks, 90% of the variance was explained by one PC, whose eigenvector will be denoted as a postural synergy. Next, based on their similarity, we clustered the obtained synergies (PCs) corresponding to the same amount of variance. In general, this led to a set of groups *Gi*, with *i* being the number of PCs at each variance level obtained separately across time windows (see Materials and Methods and Fig. S1d). In this experiment *i* = 1. To set the number of classes for the clustering, we performed the Variance Ratio Criterion (VRC) analysis (see Materials and Methods). As shown in Figure 2d (dashed line), a clear peak corresponding to two classes was evident. Thus, for Task 1, despite PCA extracting only one synergy, the successive clustering correctly splits this PC into two clusters (Fig. 2b). These seem to match the distinct underlying movements (Fig. 3a). As a quality check, the mean correlation between the eigenvectors within clusters resulted significantly stronger than across clusters. This result was obtained through a t-test (ind. samples, alpha = 0.05) (Fig. 2c).

**Figure 2.**
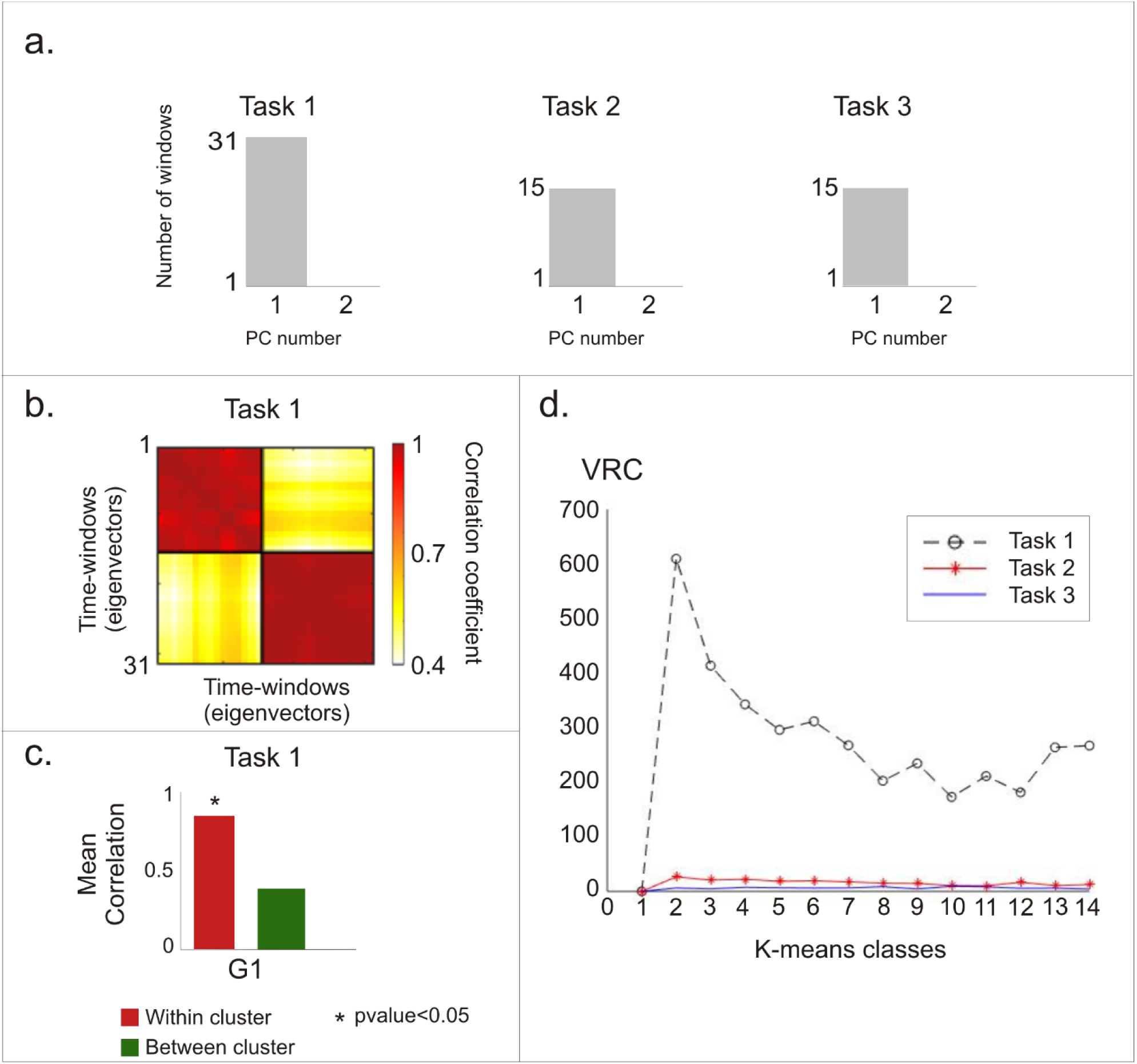
Synergies for canonical movements. (a) In the three canonical movements, PCA shows that a single PC explained 90% of the variance. (b) In Task 1, after PCA each time-window is characterized by a specific eigenvector. The first level clustering divides these eigenvectors into two different groups. Hot colors show high correlation coefficients. (c) The mean correlation within clusters is significantly higher than across clusters (t-test). Asterisks denote significant differences (p<0.05). (d) VRC analysis shows that for task 1 the optimal number of classes is 2 (grey line). For tasks 2-3, the clustering failed to separate two or more classes (red and blue line).

**Figure 3.**
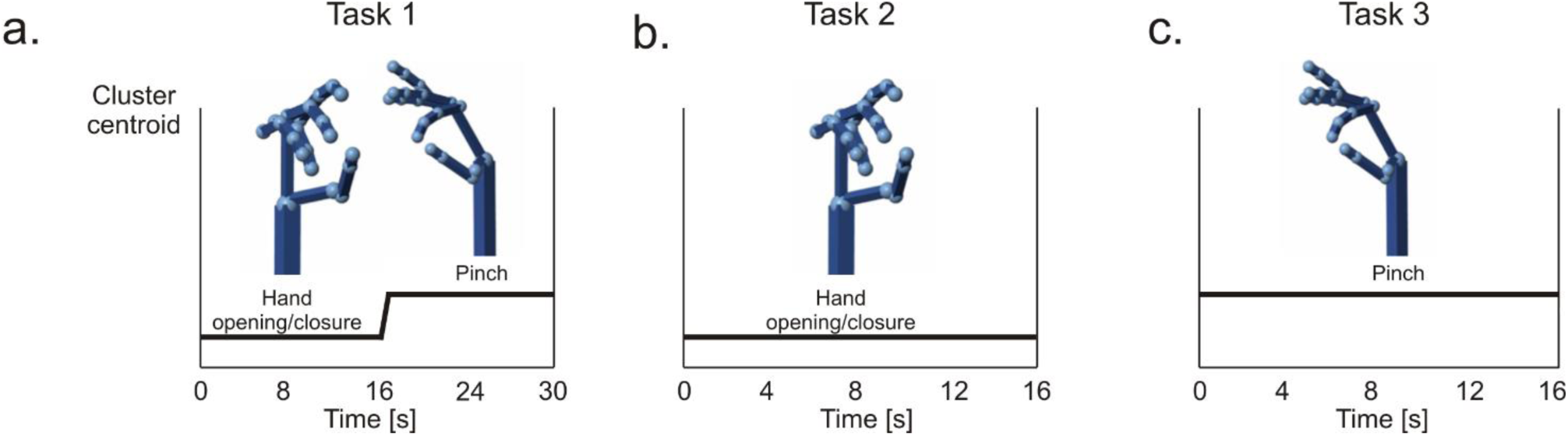
Temporal dynamics of canonical hand movements. Hand configurations were reconstructed by mapping the centroids of each cluster through a 3D Simulink hand model. (a) Task 1: sequence of hand opening-closure and pinch movement, over time. (b) Task 2: hand opening- closure movement, over time. (c) Task3: pinch movement, over time. For tasks 2 and 3, the eigenvectors belonged to the same cluster. Note the complete match between the reconstructed and the canonical/executed movement (shown in Fig. 1b).

Then, we analyzed tasks 2 and 3 as performed separately, i.e., in different recording blocks. In this case, we found, again, a single PC. However, this time, the clustering failed to identify separate clusters for these tasks. The VRC analysis (Fig. 2d) shows the absence of an optimal number of classes for the clustering (see Materials and Methods) for Tasks 2 and 3. This is encouraging: the adopted approach suggested not to split the synergy into separate groups.

These results support the idea of the low dimensionality of the hand movement. Despite a limited set of synergies explains manual behavior, different postural configurations can be linked to the same synergy.

### Temporal dynamics of canonical hand movements

Next, to better elucidate the correspondence between synergies and hand configurations we exploited a Simulink model. The 17-dimensional centroids of each cluster *Gi* were mapped to the weight of each hand joint of a 3D model using the Simulink toolbox (see Materials and Methods). This intuitive visualization represents the contribution of a synergy to the movement (Fig. S1f-g).

For task 1, the model revealed two hand configurations characterized by a precise temporal ordering, consisting of hand closure followed by the pinch (Fig. 3a). The two clusters consisted of a balanced number of time windows (n=8), resulting in a movement lasting 16 seconds (2 seconds for each window). This matches the temporal sequence of the experimental design (Fig. 1b). In contrast, a single hand movement was recognized in tasks 2 and 3 (Fig. 3b-c). These results suggest that the association between motor synergy and hand configuration is not unique. Instead, a subspace of postural synergies can explain natural hand behavior and capture its temporal dynamics.

### Natural hand movements: a complex spatial scenario of motor modules

Once the procedure was tested on the canonical movements (without object contact), we applied it to data collected in the naturalistic scenario. Here participants engaged spontaneously with everyday objects (Fig. 1a). In this case, we retrieved four groups (*Gi*, *i*=1,…,4) composed of multiple PCs explaining 90% of the variance. As before, within each group, we clustered the eigenvectors. To check the cluster quality, we report in Fig. 4a (for a representative subject) the correlation matrix of the eigenvectors ordered by cluster labeling. Please note that each eigenvector corresponds to a time window. The distance within and across clusters was tested on all subjects using a t-test (alpha = 0.05) (Fig. S2). We obtained that, across participants, the mean correlation within clusters was always significantly higher than the correlation between them (Fig. 4b).

**Figure 4.**
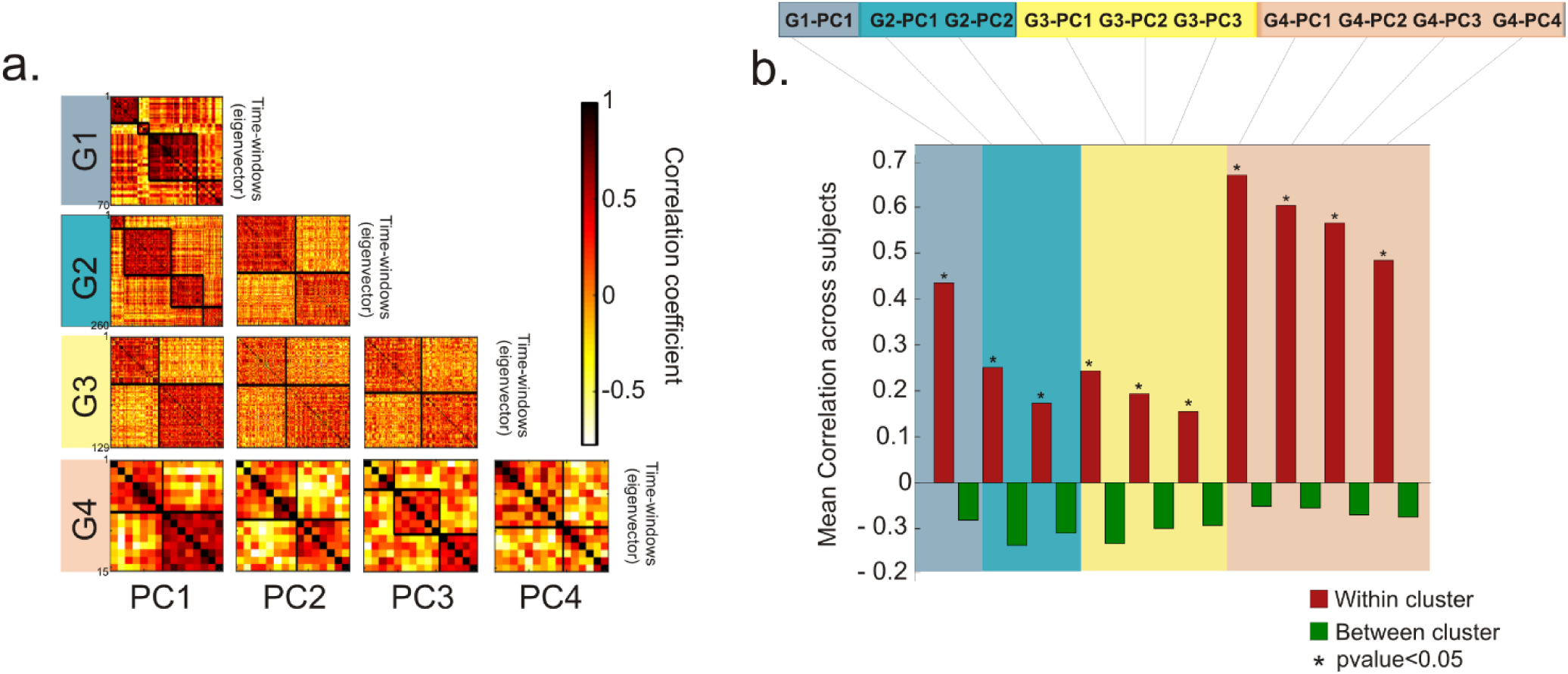
Natural hand movements: a complex spatial scenario of motor modules. (a) For a representative subject from the naturalistic task, we report the correlation matrix for each Group (*Gi*) of eigenvectors ordered based on the variance explained (each group is depicted by a different color). The blocks along the diagonal show the correlation of the eigenvectors within the same cluster. (b) Mean correlation across the thirty-six participants within (red bars) and between (green bars) clusters for each group: the mean correlation within clusters is significantly stronger than the correlation between the clusters. This effect is tested (t-test) by averaging the correlation values of all subjects. Asterisks denote significant differences (p<0.05).

Furthermore, within the same group, different clusters were extracted, e.g., in G1, four of them were identified. However, synergies might be similar across groups, i.e., independently of the amount of explained variance. This can occur if, in different time epochs, the hand configuration is either a predominant or a minor component of the movement. The first scenario will be captured by synergies in G1 (corresponding to the high variance explained). Alternatively, it can be captured at a lower level (e.g., G4 at a lower variance). This is supported by recent findings showing that synergies can manifest individually or in combination^10^. Low-variance PCs seem to reflect an interesting structure and can contribute to the precise pre-shaping of the hand to an object^18^. Therefore, it is important to group the obtained hand configuration independently of the variance explained, i.e., across *Gi*.

To identify similar synergies, we performed the agglomerative clustering on the extracted hand configurations across the different groups *Gi*. The analysis was performed separately for each participant (see Materials and Methods). Figure 5a depicts the correlation matrix of the obtained cluster centroids in four representative subjects. As expected, values were higher within- than between-clusters, in all participants (Fig. 5b). These differences were tested using a t-test (alpha =0.05). As noted in Figure 5c, only a minority of clusters were formed by centroids from the same group *Gi*. For example, in subject 7, cluster 1 was composed of two centroids from G3 and two centroids from G4. We observe that a mix of different centroids is grouped through the whole range of possible permutations (up to 6 with 4 PCs). This shows that different amalgams of motor modules come into play when the hand interacts with the external object to solve naturalistic tasks.

**Figure 5.**
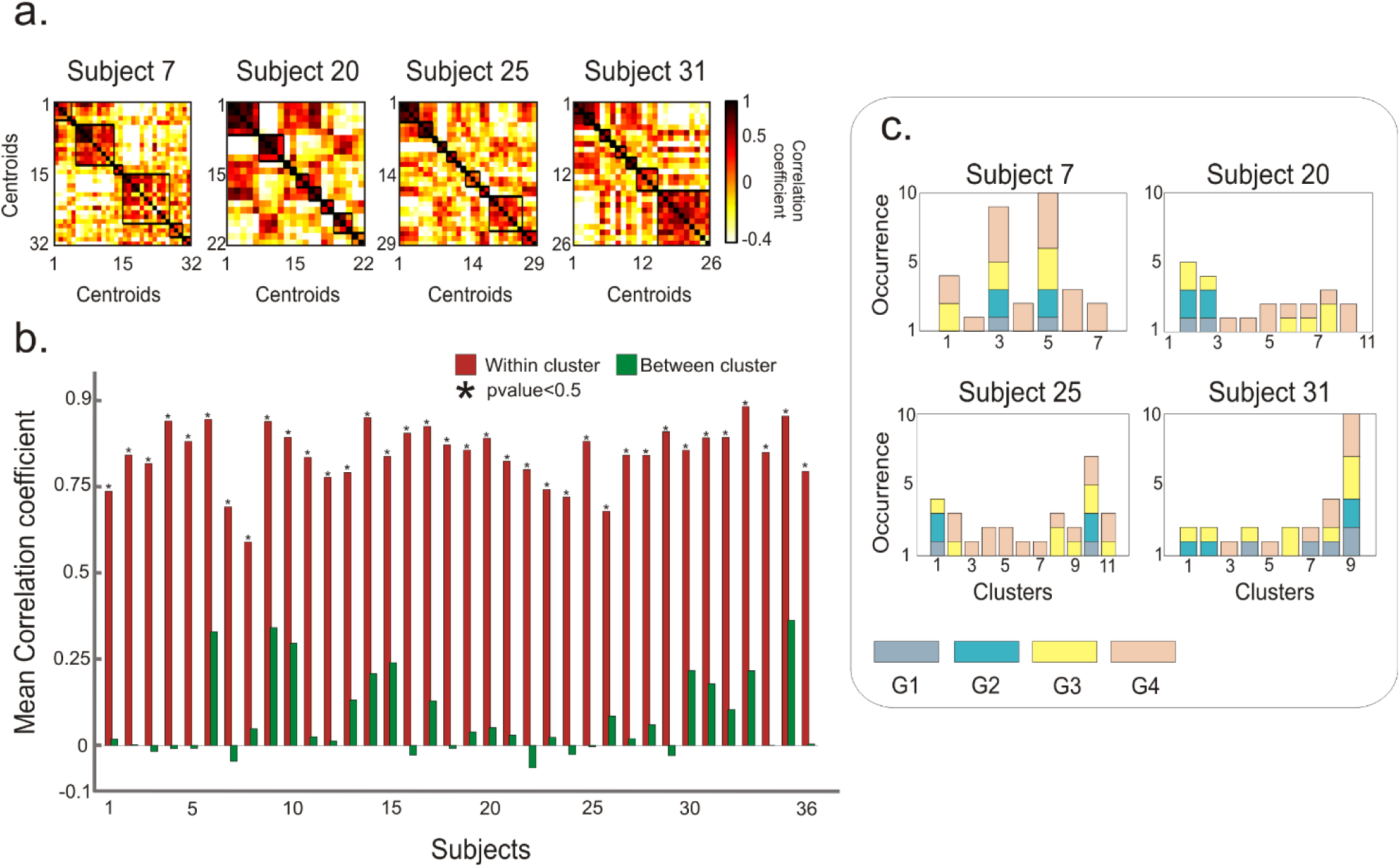
Amalgams of motor modules. We report, as an example, the results from four representative subjects. (a) Correlation matrices obtained after the second-level clustering performed on the centroids obtained from the first level clustering (shown in Fig.4). (b) Mean correlation coefficients within- (red bars) and between- (green bars) clusters, in each participant. The mean correlation within clusters is statistically stronger (* correspond to p<0.05, t-test) than the correlation between the clusters. (c) The majority of clusters comprised centroids belonging to different PC groups. The bar colors refer to the group to which the centroids belong (*Gi*). For example: in subject 7, the third cluster is composed of centroids from all 4 PC groups.

### Temporal dynamics of natural hand movements

To study the dynamics of natural hand movements, we moved into a group-level analysis by defining the hand states, *Si* (with *i*=1,…,7) (see Materials and Methods). It is noteworthy that *Si* represents centroid positions extracted from different participants, interacting with different objects. Within the same participant, they may reflect different movement phases (transport, grasp, manipulation) (see Fig. S3). In other words, they reflect prevailing motor patterns, capturing various spatial configurations of the hand. As it can be seen in Figure S3, among them, we can recognize functional movements similar to pinch grip (S1), large power grasp (S4), sphere power grasp (S5), and tripod grasp (S6). Moreover, we found the wrist planar flexion/extension (S2), ulnar deviation (S7), and the abduction of the four fingers combined with the thumb rotation (S3). Notably, these hand states resemble those found in previous studies ^10, 13, 35^.

Next, we investigated the frequency of occurrence and transitions of these hand states. Now, since the last clustering mapped each time window, of each subject, into one of the seven hand states, this provided us with a time-course of each *Si*. Hand states are mutually exclusive by definition; thus, we can define a transition from one state (starting state) to another (landing state) based on their time course. For a representative subject, the time-course of the transitions is shown in Figure 6a.

**Figure 6.**
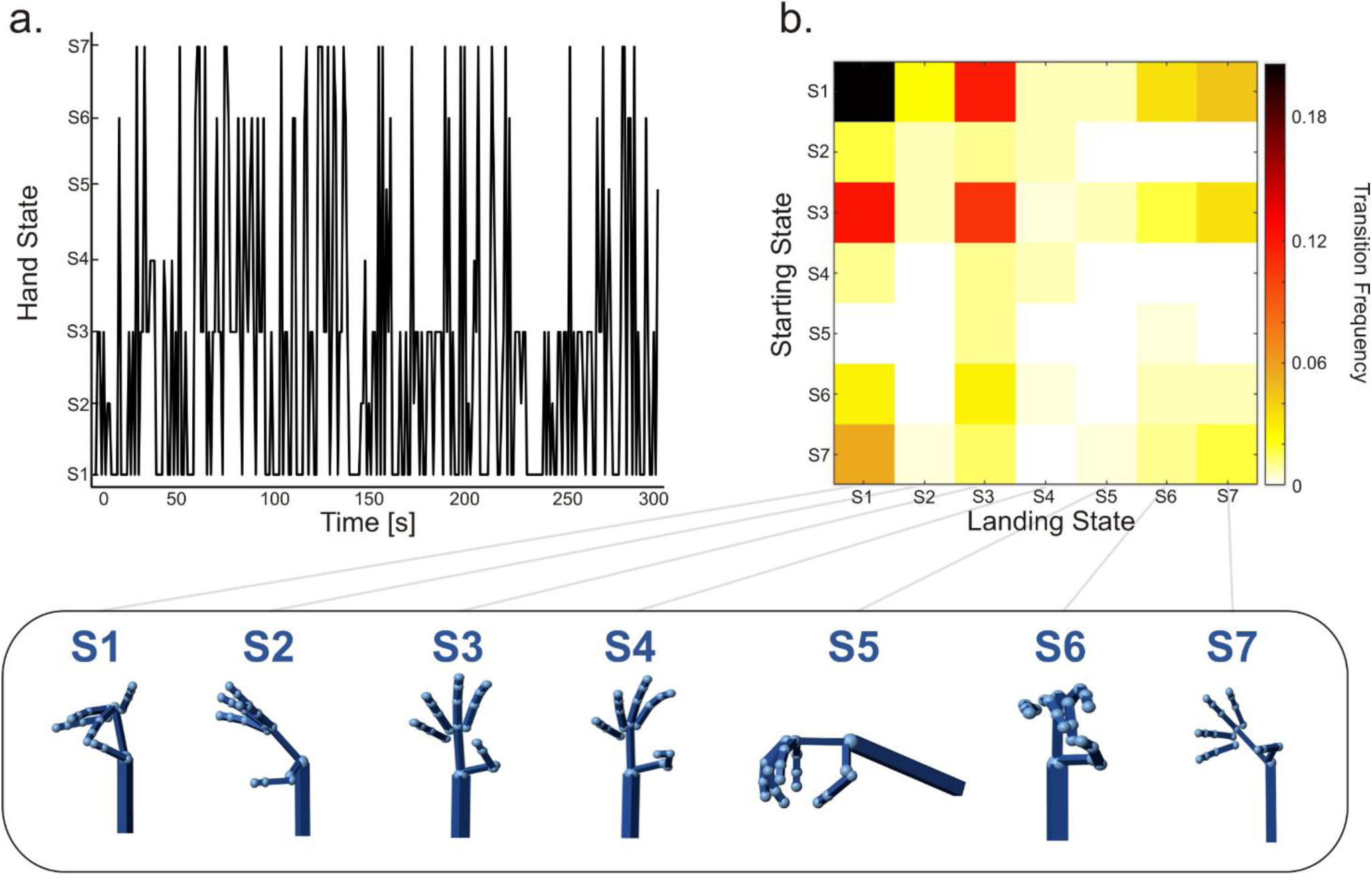
Hand state transitioning. We report the results for a representative subject. (a) Time- course of the transitions across hand states. Each transition is defined as the flow from one state (starting state) to another one (landing state). (b). The transition frequency matrix shows that some transitions are more likely to occur than others (hot colors). In this participant, the most frequent transitions are between states S1-S3, S3-S1 and S7-S1. Values along the diagonal represent the most stable states, i.e., S1 (pinch grip) and S3 (thumb rotation with the abduction of the other four fingers). These states are those in which the hand remains still without transitioning into another for a while. Below, we report the hand states mapped by the Simulink model.

It can be noted that natural hand usage continuously flows across different states. The transition frequency matrix (Fig. 6b) illustrates how often a transition between a given pair of states occurs. The frequency is computed as the number of times the transition is observed, normalized by the time of the experiment. To help the interpretation of these transitions, the seven hand states are reported below the transition matrix. For instance, from a given starting state, e.g., S1, the most recurrent landing state is the S3. This means that in this participant, the probability that the hand flows from a state representing a precision grip to a state representing thumb rotation and the abduction of the remaining four fingers is higher than for the other states. We note that the transition frequency matrix is not symmetric, e.g., the frequency of the transition S1-S6 is different from S6-S1. Further, when the hand enters a given state, it can stay there for a while (values on the diagonal). For example, S1 and S3 are also the most permanent states in this participant (see Figure 6b). Based on our findings, these hand states seem to be the most consistently recruited in using the different objects (in this subject).

Next, we investigated the consistency of the observed transitions in our sample. Figure 7a reports the percentage of participants showing a specific transition, i.e. the element (*i,j*) represents the number of subjects showing a transition from state *i* to state *j*.. For representative purposes, in Figure 7b, we report the most consistent transitions across participants and the associated hand states reconstructed from the Simulink model (see Materials and Methods).

**Figure 7.**
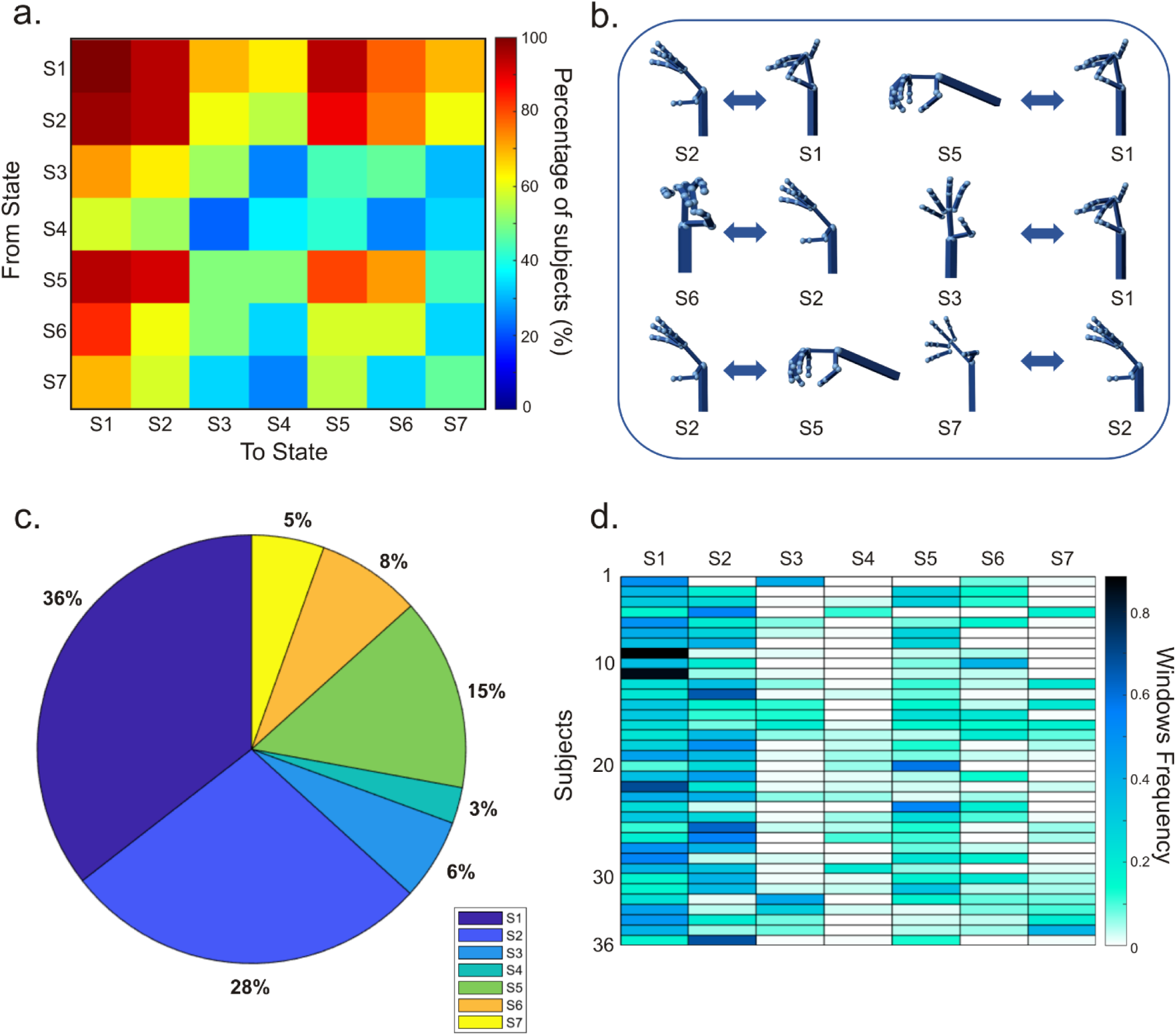
Temporal dynamics of natural hand movements. (a) The matrix shows the overall transitions observed across subjects. Each cell illustrates the percentage of participants showing a given transition. Some transitions are more consistent than others (hot colors). Note that the most likely transition is S1-S2. (b) Display of the most frequent transitions observed in panel a. The pinch grip (S1) is preceded by the flexion/extension of the wrist (S2) for 97% of the subjects. A slightly lower percentage is observed by the opposite transition (94%). S1 is also preceded by a sphere power grasp (S5), for 94% of participants. Tripod hand configuration (S6) is the landing state of S2 for 75% of participants. The opposite flow is less consistent (61% of participants). The transitioning among thumb rotation (S3) and pinch grip (S1) is found for 73% (for both directions). S2 flows to S5 for 88% of the participants and 58% of subjects show the switching behavior around the wrist movements (from S7 to S2). (c) Percentage of time spent in each state (see colorbar). For the sake of simplicity, each state is depicted by a different color. (d) For every subject, we report the number of windows associated with each state normalized by the participant’s total number of windows.

We observed high consistencies across participants for the transition S2-S1 (97%). Similar consistencies were also found for the opposite transition (S1-S2: 94%). These transitions represent the most robust strategies to achieve the hand opening/closure contingent upon the holding/releasing of small objects. It is noteworthy that S1 and S2 also show a flow from/to S5. This could represent the adaptation phase of the hand that tends to assume the optimal position for the grip. It is important to highlight how transitions involving S1 and S2 are highly frequent and consistent in most participants (Fig. 7a, first cluster along the diagonal). These two states seem to represent central hand shapes attracting all the other ones. This suggests that the flexion of the wrist and the closing of the middle finger towards the thumb represent two essential motor patterns in the reach-to-grasp movements.

Furthermore, if we consider the tripod hand configuration (S6) as the landing state, the flow from S2 is more consistent across participants than the opposite flow (S2->S6:75%; S6->S2:61%). This transition may suggest that the flexion-extension of the wrist is most plausible as the first stage of the movement before contacting the object. Once the object is grasped, it is reasonable to assume that the wrist remains stable. Remarkably, the alternance of S6-S1 is higher than the stability of S6. This transition is a physiologically plausible representation of the transition between two precision grips to move sequentially across different objects. This sequence might be due to the naturalistic context of our study. More than 50% of participants used thumb rotation (S3) to flow towards S1 and S2, while less than 50% of the participants showed transitions to other states. Regarding the switching behavior around the wrist movements (states S2-S7), the transitions involving S2 are more robust than those recruiting S7 (on average, 73% vs 46% of participants, respectively). This suggests that reach-to-grasp movements are more associated with flexion-extension of the wrist (both before and after grasping the object) than with ulnar/medial deviation movement.

Figure 7c reports the overall proportion of windows assigned to every *Si*. This shows how often a state is involved with the movement. Interestingly, three states, S1, S2, and S5, jointly account for approximately 80% of the motor strategies chosen by participants to solve the naturalistic task. Notably, this set of states largely recapitulates the commonly engaged hand movements, i.e., precision grip (S1), spherical grip (S5), and wrist flexion (S2). The remaining states seem to play a minor role occurring around 20% of the time. Within different participants, hand states show different frequencies of occurrence (Fig. 7d). The matrix reports the number of windows associated with state *Si* normalized by the total number of windows of each participant. Finally, in Figure 7d, we report the frequency of occurrence of the states for each participant (see Materials and Methods). S1 and S2 are the most observed states in all 36 participants, followed by S5 and S6. The remaining states occur less consistently across subjects.

In conclusion, we characterized natural hand movements as a switching mechanism between a discrete set of hand states. The analysis of the most consistent transitions across participants shows some regularity of interaction with everyday objects. Such regularity is notable since participants performed hand movements in an unconstrained way.

## Discussion

This work examined how natural hand movement unfolds in space and time in a continuous flow of everyday acts. We modeled the architecture of manual behavior as a sequence of hand states. We inferred the state’s spatial structure and temporal dynamics. We hypothesized the presence of regularities, both in space and time, even in a naturalistic task. We developed a combined approach based on PCA and clustering techniques to test this hypothesis. Once validated in a controlled experiment, we applied it to natural movements. Our findings show a complex spatial organization of hand states built from amalgams of basic motor patterns alternating over time through specific transitions.

### The functional role of motor synergies

Previous studies report that the hand movement lies in a limited dimensional space, spanned by canonical synergies. This was shown in laboratory-controlled experiments ^10, 13, 28^, and in everyday manual behavior ^27, 36^. We identified motor synergies in line with this hypothesis, but the spatial scenario was more complex than expected. Two important features emerged. First, synergies explaining the same amount of the variance corresponded to different hand movements: in different temporal windows, distinct motor patterns can play the same role (Fig. 4). Second, in different time epochs, the same hand configuration contributed to a different degree, i.e., with a major or a minor role, to the movement observed (Fig. 5). Thus, the association between motor synergy and hand configuration is not unique.

In line with these findings, synergies underlying different movements have been reported in muscle activations of freely moving animals ^21, 37^. This implies that the explained variance cannot be the only leading criterion for extracting motor primitives. In the construction of the natural movement, very similar covariation patterns subserve very different hand configurations. This also applies to low-variance PCs. Classically considered as noise (in motor terms), more recently, they have been described as structured and task-related^18^. Of note, we found that, in specific time instances, PCs explaining small and high variance often share the same spatial structure (Fig. 5). This structure drives the identification of the hand states (Fig. 6, S3). Therefore, multiple configurations underlying the same synergy drive the functional significance of the natural movement. For example, the canonical experiment shows that the extracted synergy cannot disentangle the two underlying functional movements captured by the postures.

This finding provides insights into the role of motor synergies. One leading idea is that kinematic or muscle synergies reflect anatomical/biomechanical constraints^2^. This is supported by observations that not all the hand joint angles are controlled independently of each other ^10, 27^. Even when subjects are instructed to move one finger, the correlated motion may occur in adjacent fingers^38^. Nevertheless, a motor synergy is also pluripotent, as it contains information on the spatial structure of the fingers, allowing the emergence of hand configurations adapted to the environmental demands. The object’s properties dictate these variations to achieve the sophistication inherent to dexterous manipulation.

### The spatio-temporal architecture of hand states

Behavior does not exist in isolation, instead it is often described as organized in chains (e.g., the morning routine to preparing breakfast). Analogously, we modeled manual behavior as shaped by sequences of simple motor acts building complex movements. We characterized the temporal dynamics in terms of the occurrence of hand states and their switching (defined as transition frequency). Similar approaches have been successfully applied to characterize the transition across emotional states^39^.

We identified a set of hand configurations consisting of wrist movements (i.e., S2, S7) and metacarpophalangeal joints of the thumb and fingers (i.e., all other states) (Fig. 6, S3). Some of these hand states may relate to different functions^40^. For example, the involvement of pads of the digits (S1, S6) recalls a precision grip, whereas all fingers opposing the thumb (S4, S5) remind whole hand prehension. A precision grip typically involves the index finger opposing the thumb. In our case, instead, it consists of more than one finger opposing the thumb. We already know that both intrinsic (physical features) and extrinsic properties (location in space) of the objects influence the grasping kinematics and hand shaping^41–43^. Small objects are preferentially grasped through a precision grip^41^, but depending on shape, size, and orientation, individuals prefer recruiting more fingers to grasp them stably^43^. In our study, objects have different shapes (characterized by size, shape, and diameter, see Table S1). Furthermore, their relative orientation dynamically varies depending on how participants grasp and act. Thus, compared to previous studies, where movements were highly controlled and reproducible^10^, here the extracted hand states encompass a larger repertoire of motor adaptations. These will be contingent upon intrinsic and extrinsic properties, and different objects’ handling might contribute to the same state. Thus, unlike^10^, where movements were confined to a single imagined object, in our case, a state cannot be uniquely associated with an object. Different objects and individual motor patterns contribute to the extracted state. This may explain why, for example, in the precision grip state, we observed deviations from the expected classical grip structure.

Apart from the spatial organization of hand states, their temporal transitions play a fundamental role in understanding how participants construct everyday movements. Most transitions displayed high inter-individual stability, i.e., found in almost all participants. These transitions could not be predicted by biomechanical constraints^2^. They reflect alternation of hand configurations with specific functional roles. Beyond actual grasping, we recognize hand states resembling the first phase of grip formation (S3). We identified this phase through specific trajectories of hand state transitions. Classically, the kinematics of grasping consists of two main phases. A progressive grip opening with straightening of the fingers (transport component) is typically followed by a gradual closure^44^. In our data, the alternance between S3 and grasp-related hand states (S1, S5, S6) may resemble such kinematics (Fig. 7a). However, in the majority of participants (72.2%), the most common landing state is S1. The inverse transition (S1-S3) was slightly less consistent across participants (69.4 %). One possible explanation is that, once the object is grasped, the hand more frequently flows through states enabling prehensile acts. This is supported by the high consistency observed in transitions across grasp types (from/to S1, S5, S6) and participants. Plausibly, this might depend on our task paradigm.

Here, participants were asked to arrange all the objects on the table and then have breakfast. Accordingly, we expect the transport component of the hand from its initial position to the target to be especially involved in the first phase of the task (i.e., when the hand needs to reach objects located at different distances). Once the table was set, all the objects were in the participant’s proximity. This condition encourages a direct switch between hand states associated with the object manipulation, skipping the necessity of transporting the hand over long distances. The second most common hand state is the wrist flexion-extension (S2) (Fig. 7c). We know that this pattern belongs to a plethora of movements employed in the interaction with objects^16^: accordingly, the high consistency is observed in almost all subjects and time windows (Fig. 7d). The transitions with the other grasp-related states (S1, S5, S6) demonstrate that S2 is instrumental to the grasp construction (Fig. 7a). By contrast, the ulnar deviation (S7) seems less involved in our task.

Once the hand enters a specific state, it often remains in the same position without transitioning into another for a while (Figure 6-7, values along the diagonal). We speculate that this regularity corresponds to an efficient strategy of optimization costs to maintain fingers in an unaltered posture. This was common across subjects and it aimed at stabilizing hand-held objects. Specific hand states (e.g., S1, S2, S5, S6) were consistently stable in more than 50% of participants. Of note, our method cannot discern if the observed hand configuration was engaged in a movement or the hand idles ^27^ .

To summarize, the observed transitions are not randomly distributed. Certain motor patterns are likely to occur after others, thus defining specific trajectories in the hand state space. These were highly consistent across subjects. Of note, we observed a few possible transitions among hand states, able to explain a large variety of movements. We speculate that this corresponds to a temporal mechanism that simplifies hand motor control. This temporal feature integrates with the spatial features discussed above, characterized by a large degree of complexity due to the entanglement between synergies, hand configurations, and final hand states. Thus, based on our findings, and differently from previous works ^10, 13, 16^, we suggest that the simplification of the motor control unravels across the temporal dimension more than the spatial one. These findings open new perspectives on how these temporal structures are mapped on the neural dynamics.

## Conclusions

Natural hand movements in naturalistic settings show regularities in space (structure) and time (transitions). At the spatial level, these regularities do not necessarily solve the problem of simplifying the control strategy of the hand. By contrast, this might take place in a temporal dimension. This is realized by reducing the number of transitions among a finite number of hand states. We expect the experimental contexts or pathological conditions to modulate such spatiotemporal structure tuned to specific temporal features of the movement itself, similar to what is found for the structure of synergies^36^. In conclusion, our study paves the way for future investigations on neurophysiological correlates underlying such simplified motor strategies.

## Materials and Methods

### Experimental procedure

Thirty-six healthy participants (mean age 26.61, females 16, males 20) performed reach-to-grasp movements while interacting with everyday objects aimed at preparing and having breakfast. The objects used ^10^ were: a placemat, a napkin, a plate, a fork, a knife, a spoon, a glass, a bottle of water, a jug, a tea bag, a coffee cup, a cup, a sachet of sugar, a teaspoon, a circular ashtray, a packet of cigarettes, a toothpick, a jar, a frying pan, an egg and a salt container. The height, weight and diameter (or width) of each object is reported in Tab S1. Subjects were asked to sit in front of a table where objects were arranged (Fig. 1b) and interact with them in a naturalistic way. The instruction was to prepare and have breakfast using all available items. Finally, they were asked to clean and set the table. Participants gave their written informed consent. The procedures were approved by the IRCCS Fondazione Santa Lucia ethics committee and were in accordance with the standards of the 1964 Declaration of Helsinki.

### Data acquisition and calibration

For each subject, kinematic data were acquired by means of a right-handed, 18-sensor CyberGlove (CyberGlove II System, Virtual Technology, Palo Alto) equipped with two band sensors per finger, the metacarpo-phalangeal joint (MCP) and the proximal interphalangeal joint (PIP), and four abduction sensors, thumb crossover sensors, palm arch, and two wrist sensors (flexion and abduction). Since the palm arch sensor was set to zero (due to technical problems), 17-sensors were used to acquire the data. The CyberGlove is designed for tracking hand kinematics, by measuring joint angles with resistive bending sensors. Once calibrated, the voltage signal related to the joint-angle data was digitized. The instrument was connected to a host workstation via bluetooth through a serial port. To establish and manage this type of communication, a MATLAB interface was developed. For every participant, markers positioned at fiduciary points were applied to the glove for the calibration step.

To maximize the accuracy of calibration, a two-step procedure was developed. First, the participant was asked to keep the right hand still in a specific reference hand posture (Tab S2). A picture of the hand was taken for each posture and, at the same time, the output values of the 17 sensors of CyberGlove were recorded. To evaluate the joint angles from the acquired pictures, we used a binocular stereophotogrammetric system consisting of two GoPro Hero3 cameras (https://gopro.com/it/it/update/hero3) mounted on a common rigid support. Intrinsic and extrinsic calibration parameters of the stereo-system were computed by using a MATLAB Camera Calibration Toolbox (https://doi.org/10.22002/D1.20164). Thus, through the spatial positions of fiduciary points, we were able to measure the joint angles from the previously acquired pictures. We developed a MATLAB interface to extract 17 different angular values from 10 reference postures of the hand (Table S1). This allowed us to overcome the inter-subject anatomical differences, and improve the reproducibility of the measurements. For each joint, we selected two images where the joint presented the minimum and maximum angle and we extracted the three points referring to the joint angle (by using the markers previously applied to the glove). The second point was the vertex of the angle to be estimated. The final step of the calibration consisted in computing the gain and offset parameters^45^. To his aim, a linear law was employed, as follows. For each sensor, two pairs of raw values of voltage (mV) and angle (degrees) were acquired. The calibration law is, therefore, defined by the system of equations:

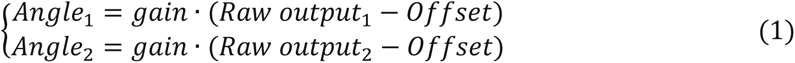

where Raw output1 and Raw output2 are the raw voltage values registered by the sensor, Angle1 and Angle2 are measured angles associated with the sensor.

Thus, gain and offset can be computed by solving the linear system:

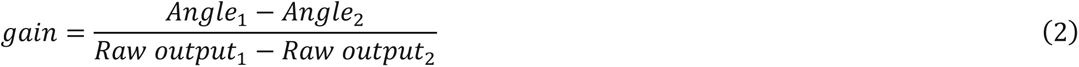

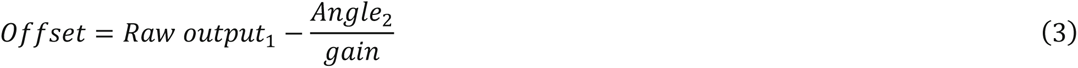

under the hypothesis that *gain* ≠ *0* or, equivalently, *Angle*_1_ ≠ *Angle*_2_.

At this point we were able, for any raw output of a sensor, to compute the corresponding joint angle.

### Data analysis

This paper tests the idea that natural hand usage can be modeled as a discrete set of hand states alternating with a precise temporal order. To extract the final hand states from the acquired data is a complex and challenging task. For this reason, our analysis is driven by several assumptions, built through previous experimental evidence. First, for every subject, we assume that a large amount of data variability can be explained through a few motor synergies or patterns (e.g., ^10, 18^). Second, as we employ a naturalistic paradigm, we expect hand movements to vary over time as a function of the phase being considered (e.g., hand transport or object contact)^17^. These changes may arise from a specific combination of the same set of synergies which thus might play different roles in different temporal epochs. For example, in a given time window we might observe one main movement captured by a single synergy corresponding to a large amount of variance explained. On the other hand, in a different temporal instance, we might observe a complex movement that is captured through a larger number of synergies. In this case, each of them will explain less variance than before. Previous evidence supports this idea by showing that the gradual molding of the hand occurs in a two-dimensional space, in which the first two PCs can manifest individually or in combination^10^. Thus, in different time windows, the same pattern might have a large/small weight and corresponding explained variance, if it represents a main/minor component of the observed movement. Third, we expect, as in.^26^, these synergies to be reproducible at the subject level. Thus, we group them by adopting a two-step procedure. First, we cluster synergies separately for the different levels of variance explained. Then, we group them across the different levels of explained variance. This corresponds to putting together those synergies that play different roles in different time windows, at the subject level. Eventually, we cluster them across subjects to obtain the final hand states.

Based on these assumptions, our analysis can be summarized as follows (Fig. S1). First, for every participant we extract for every temporal window (2s), the most recurrent synergies through PCA (*dimensionality reduction)*. Second, through a K-means at each level of variance explained, we cluster synergies with similar structure *(first level clustering)*. Third, we group together the obtained synergies’ centroids across the different levels (*second level clustering*). Fourth, a final K-means clustering was performed on the individual patterns, across subjects (*group-level hand states).* Finally, we investigate the temporal dynamics and transitions among the obtained hand states (*hand states dynamics and transitions)*.

### Dimensionality reduction

To identify the most recurrent patterns in the acquired data, we adopted PCA. It consists of a linear reduction dimensionality technique that provides a parsimonious representation of hand kinematics into a set of principal components (PCs) (or eigenvectors) ordered according to the percentage of variance explained (eigenvalues)^14, 18^. Each eigenvector leads to a hand configuration, since each vector component represents the weight for a specific hand joint. The kinematic data were filtered with a low pass FIR filter (passband frequency = 2 Hz and stopband frequency = 3 Hz). To investigate the dynamics of the naturalistic hand movements, signals were sampled with overlapping windows (windows length = 2s, overlap = 50%). In every window, PCA was computed on sensors’ time course to extract principal components (PCs) accounting for at least 90% of the observed variance. The length of the window was chosen to guarantee a good compromise between the spectral content of the signal (no significant contribution was observed beyond 0.25 Hz, not shown), the time resolution and the number of samples needed to get stable estimates of the PCA eigenvectors and a good ratio between the number of temporal samples (n=40) and variables (n=17, number of sensors).

After PCA, every window is characterized by a number of eigenvectors based on which the windows have been divided into 4 groups: group 1 includes windows with only one eigenvector (i.e., only one PC explains 90% of the data variance), group 2 includes windows with two eigenvectors, group 3 windows with three eigenvectors and group 4 windows with four or more eigenvectors. Within each group the eigenvectors are ordered based on the variance explained, i.e., the first eigenvector explains more variance than the second and so on.

Based on our hypothesis, we expect the number of eigenvectors to relate to the weight of each synergy on the movement performed. For example, we expect a time window characterized by a main movement to be characterized by a single PC with the largest weight (falling in G1). On the other hand, G4 will relate to movements that are explained through the composition of more than 4 PCs each of them with a small weight (G4).

Of note, at this stage, since the analysis is driven only by the variance in the data, the hand configuration underlying the synergies does not play any role. Two eigenvectors explaining the same variance, extracted from different time windows, may relate to completely different hand configurations. For this reason, once extracted, we proceed to cluster the obtained PCs based on their similarity.

### First level clustering

Now, we clustered the obtained PCs ranked by the amount of variance explained. We assume that depending on the movement performed, we might identify different hand configurations underlying the same PC (or synergy). This is because the amount of variance explained by a given PC may depend on the predominance of this synergy within the movement. One PC (or synergy) suggests a single main movement. However, this movement can be different in different time windows. Thus, within each PC a certain number of distinct hand configurations can be identified.

We considered the first group (G1) composed of patterns with the highest weight (the main movement explained by one single eigenvector) and the lowest group (G4) composed of patterns with the smallest weight (the main movement explained by a larger number of PCs, namely more than four).

For each participant in a given group *Gi* (*i=1,…,4*), we adopted a K-means algorithm to cluster the obtained PCs. Within each group, eigenvectors were clustered separately based on the amount of variance they explained: we cluster all the first eigenvectors (highest variance explained - highest weight in explaining the movement), then all the second eigenvectors (lower explained variance - lower weight in the movement) and so on until the last ones. For any group *Gi* we run *i* K-means independently. Based on our data, this leads to 10 K-means in total.

Specifically, in K-means we adopted the cosine distance (defined as 1- the cosine of the angle between vectors) to compute the distance between an eigenvector and each centroid. This choice corresponds to aggregate eigenvectors according to their angular proximity and not in terms of amplitude. To estimate the number of classes, *k_opt,* to be provided to the algorithm, K-means was run corresponding to different values of *k* sampled in a range that goes from 2 to the integer number closest to 95% of observations.

The optimization criterion for the identification of the optimal *k_opt* was the Calinski-Harabasz Criterion, also known as Variance Ratio Criterion (VRC)^46^. This is based on the ratio of between-clusters dispersion (SSB) and of inter-cluster dispersion (SSW) for all clusters defined as:

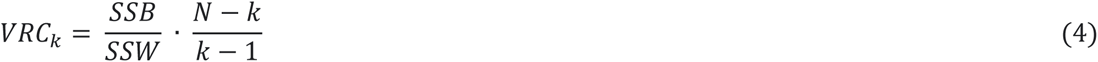

where *N* is the number of eigenvectors to cluster and *k* is the number of clusters for the current iteration. The optimal *k_opt* is obtained as the number of clusters *k* maximizing VRC.

Once *k_opt* has been estimated, we performed an independent quality check of the obtained clusters. Figure S2a shows the distribution of the correlation values among all the obtained clusters. These were statistically tested to assess the significance between the intra- and inter-cluster correlation. For each cluster, we converted the correlation value to *t*-value and, separately for every clustering algorithm, we performed a *t*-test (alpha level = 0.05). The above steps were repeated for all the groups.

### Second level clustering

Once we identified, for every subject, the most consistent motor patterns, we investigated whether some of these were similar across the different levels of variance. In fact, based on the hypotheses described above, we might have, in different windows, the same hand configuration with a different weight being involved in different phases of the movement. Now, the aim is to group together the obtained synergies across the obtained groups.

Here, the centroids obtained at the previous stage were clustered according to a hierarchical algorithm. The cosine metric was adopted for the distance and the Unweighted Average Distance was used as the cluster linkage criterion ^47^. These two metrics were adopted since they resulted in an optimal combination based on the Cophenet coefficient. The Dendrogram cut-off was evaluated according to the Inconsistency Criterion obtaining a threshold around 1 for all subjects. Also in this case, to validate the obtained clusters, we computed the average intra and inter-cluster correlation coefficient and the difference between them were tested with a t-test (alpha level 0.05). The final step of this analysis was to assign a single hand configuration to each time window.

To this aim, we proceeded as follows. So far, in every time window, we had a set of eigenvectors corresponding to the extracted PCs (from 1 to 4) which have been assigned to specific clusters. Thus, apart from the case when only one eigenvector was found (G1), any window corresponding to a set of eigenvectors mapped into *n* clusters, could not be uniquely assigned to a single cluster. For this reason, we assumed that, in any window, the hand configuration can be represented by a composite functional structure obtained by averaging the centroids of the involved clusters. For example, in a time window of 4 eigenvectors partitioned in 3 clusters, the hand configuration will be defined as the average of the centroids of the 3 clusters. In this way, each temporal window is associated with one functional hand structure.

### Group level hand states

The final step of our analysis is to map hand shapes obtained at the subject level into a set of common ones defined at the group level. To this aim, a K-means clustering across subjects was applied to previous outputs. This procedure will assign a single hand state to every time window, thus defining their temporal dynamics. The correlation was adopted as the distance metric. The number of iterations of K-means was optimized by repeating the clustering at an increasing number of iterations, until convergence was obtained. We observed the stable results when we moved from 500 and 1000 iterations. For this reason, 500 iterations were used in what follows. At every iteration, a new initial position of the cluster centroid was used. To optimize the final number of clusters, we used a multivariate index of performance^48^ based on various metrics: the average cluster size, the Dunn Index^49^ and Davies-Bouldin index^50^. K-means was performed for different values of *k* (from 2 to 25) and the optimal number of clusters *k* was established considering the stability of the performance index according to its relative maxima. We considered 3 different clustering configurations for 4, 7 and 10 classes (Fig. S3). These synthetic hand shapes will be defined as “hand states”, modeled as vectors of the same size as the PC eigenvectors. We mapped the 17-dimensional centroids corresponding to 4, 7 and 10 classes. Figure S3 shows a hierarchical scheme of the hand states: the first level of the hierarchy provides the first set of hand configurations reflecting an initial set of macro-states. Notably, they seem to characterize the primary hand motor patterns to perform in naturalistic tasks. At the second level we obtained 3 additional sub-states. This framework expands the set of functional hand shapes in everyday movements. This includes: pinch achieved through closing the middle finger on the thumb, wrist flexion, thumb rotation, index pinch, spherical grip^51^, hand closure and ulnar deviation of wrist. Now, since each hand state is represented by a single centroid, it is important to stress that the set of hand configurations falling in a given class, might exhibit a certain excursion from the average. As an example, in Figure S4 we report the first six observations closer to the mean. The final step was to map the centroids corresponding to 10 classes. The additional states found in this new configuration are either a repetition of existing states (S8 - previous S4) or noisy, i.e., joint modulations resulting in non-functional (behaviourally irrelevant) shapes (S9, S10). Notably, the obtained centroids are not driven by any a priori knowledge or constraint, i.e., they capture common movement patterns completely driven by the grasping tasks performed by all participants.

Finally, we reconstructed the dynamics of the states and computed the frequency of occurrence. The dynamics are represented through binary time courses sampled with overlapping windows (overlap= 50%, temporal resolution 1s) associating to each window its state for each of the 36 subjects (Fig. 6a). Then, the frequency matrix was evaluated by calculating the number of windows associated with each hand state and dividing by the time of the experiment. Figure 6b shows the percentage of time in which the transition between the states y (row) and x (column) occurs and how the hand continuously flows across different states. Along the diagonal (x=y) can be observed the permanence in a state.

As it can be seen in Fig. S3, S1 was characterized by the flexion of the metacarpophalangeal joints (mcp) with different degrees of flexion of the middle and ring fingers. This hand shape resembles a pinch movement, but realizes a larger excursion of the middle finger, as compared to the ring finger, that meets the thumb. Since here the index finger shows an atypical position, we showed the range of hand configurations spanned in this class, by providing the first six most correlated observations with the centroid (Fig. S3). State S2 was the planar wrist flexion/extension achieved through rotation at the radiocarpal and midcarpal joints. State S3 has predominant involvement of the thumb metacarpophalangeal rotation joint and the abduction of the other four fingers typically observed during the transport phase of the movement ^33^. S4 resembles a whole hand aperture/closure with the abduction of the thumb, similar to a large power grasp. State S5 consists of a sphere power grasp, which actively engages all fingers of the hand in order to grasp an object (e.g.^35^). The fingers and thumb flex around the edges of the object. The relative flexion of the fingers will depend on the object’s size, e.g., for a small ball these will result completely flexed. For example, participants can adopt this configuration to open the lid of the jar or water bottle and grasp the egg during breakfast. S6 shows a second precision movement, similar to a tripod grasp^35^. State 7 consists of the ulnar deviation of the wrist that allows the hand to rotate outward in the range [0°, 45°].

### Simulink model

To visualize the identified states, we built a Simulink hand model using the Simulink “Simscape Multibody” toolbox. It provides a multi-element simulation environment for 3D mechanical systems. The implemented model has the same number of degrees of freedom of the CyberGlove (17 degrees of freedom, d.o.f.). For visualization purposes, we add 5 d.o.f. that are missing on the CyberGlove, that is, a palmar arch (set to zero) and 4 sensors for the distal interphalangeal joints (DIP) of the index, middle, ring and little finger. Each finger was schematized as a kinematic chain that has its reference base in the wrist and the final effector in the fingertip. Compared to the other fingers, the thumb presents some differences in terms of joints and possible movements. The fingers, from second to fifth, were schematized with the proximal interphalangeal joint (PIP), with one degree of freedom each (flexion/extension), and the metacarpophalangeal joint (MCP), with two (flexion/extension and abduction/adduction). The thumb instead was modeled with the metacarpal-phalangeal (MCP) joint, with 3 degrees of degrees of freedom (flexion/extension, abduction/adduction and rotation around the palm), and the proximal interphalangeal joint with one. Measurements of the two wrist sensors of the CyberGlove (wrist pitch and wrist yaw) were modeled with a single joint with two degrees of freedom. In the adopted model, the segment representing the middle finger was fixed along the line passing through the MCP and the wrist joint. From this segment, the abductions of the other fingers were evaluated, excepting the thumb. Then, for each finger, the flexion and the extension of the DIP joints was assumed to be 1/3 of the flexion/extension of the PIP joints. Finally, each model component was composed of a “connection frame”: a reference system associated with each component with respect to which the geometric transformation of rotation and/or translation was performed. Eventually, we computed an artificial matrix with a range excursion for each sensor (joint of hand). We set the fingers range from 1° (minimum value) to 100° (maximum value) and the wrist range from 1° to 45°. This is in line with the natural excursion values of the hand^52, 53^. We can visualize a specific hand movement by multiplying the synthetic hand configuration (1x17 weight vector) with the base artificial matrix. Of note, a null weights vector results in an open hand posture with absence of abduction between the joints. In this way the resulting hand movement is guided only by the modulation of the joints of the synthetic posture.

## Acknowledgment

This work was supported by the European Research Council (G.A. No 759651 to VB). We thank Luca Compagnucci for his support to the software development, Francesca Leone and Alessandro Moscatelli for their useful comments on the manuscript.

## Author Contributions

V.B., C.D.G., A.P. and M.S. designed research; C.D.G., A.P. and M.S. collected the data; V.B., C.D.G, A.P., M.S. and F.d.P., contributed new agents/analytic tools; D.S., C.D.G., A.P. analyzed data; V.B., D.S., C.D.G., F.d.P. wrote the paper.

## Supplementary Information

**Figure S1.**
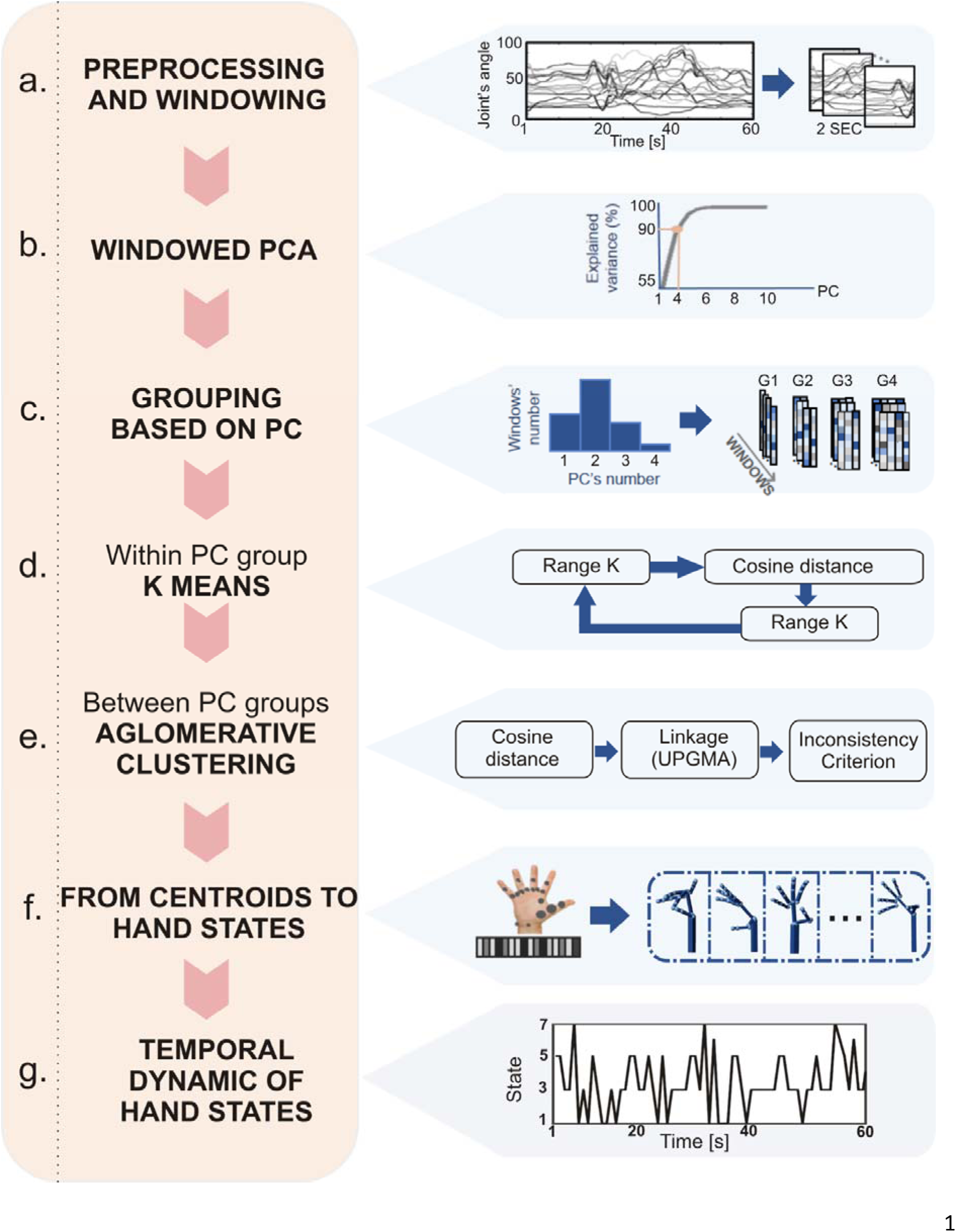
Analysis pipeline. (a) Preprocessing and windowing: the acquired kinematic data are interpolated and calibrated. A low pass FIR filter is applied, and the signals are sampled with overlapping windows. (b) Hand kinematics has a low dimensionality: for each participant, in every window, PCA is applied to extract components accounting for at least 90% of the observed variance. A number of 4 PCs explained at least 90% of the variance. (c) Group analysis across all subjects: the histogram shows for all subjects 4 PCs are enough to explain at least 90% of the variance. On the right panel, we report for each window the eigenvectors that form the 4 PC groups for a representative participant. (d) First level of clustering: K-means clustering algorithm within PC groups across the time-windows. The optimal number of classes is provided by VRC (see Materials and Methods). (e) Second level of clustering: hierarchical cluster analysis on all the centroids obtained from the K-means. (f) Final K-means clustering for the hand states: centroids of all participants were clustered to obtain a common set of hand shapes. Each 17-dimensional final centroid was mapped with a Simulink hand model (g) Temporal dynamics of hand states.

**Figure S2.**
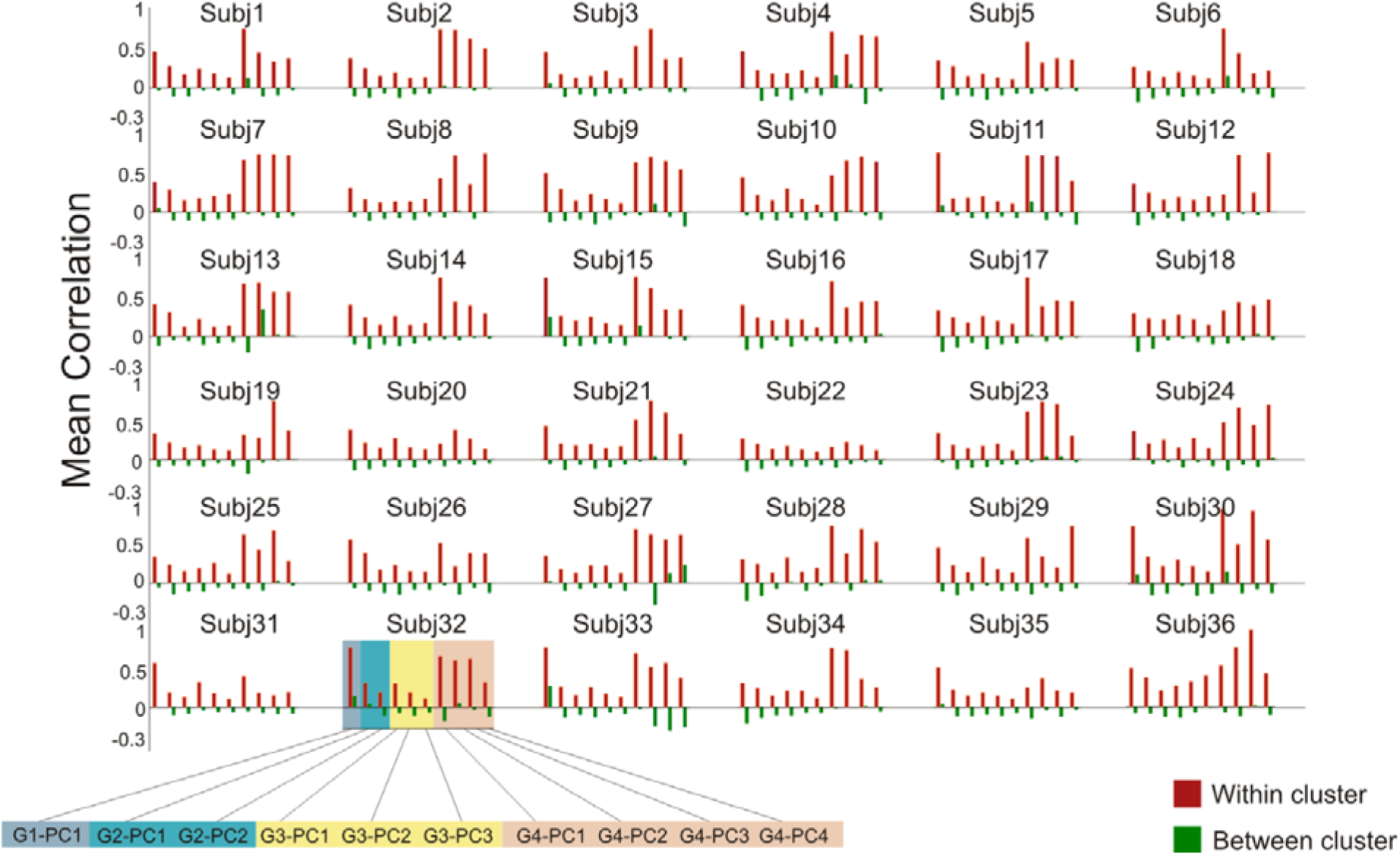
Mean correlation within and between clusters for each group and participant. The mean correlation within clusters (red bars) is significantly stronger than the correlation between the clusters (green bars). This effect is tested on all subjects using a t-test (α=0.05)

**Figure S3.**
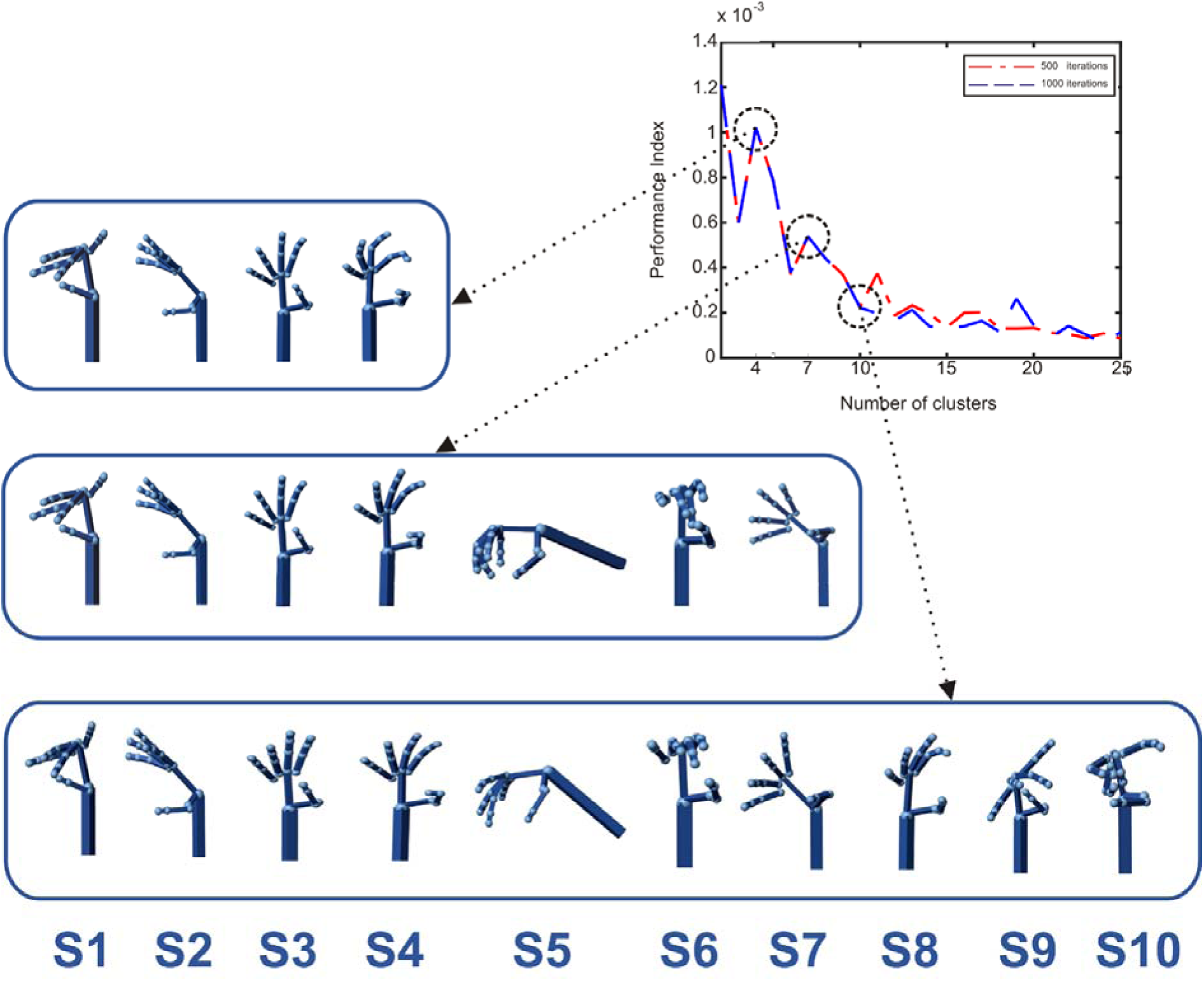
Performance index analysis to assess the optimal number of classes. The performance index as a function of the number of clusters is reported. On the left, we report different hand states obtained at 4 (first row), 7 (second row) and 10 (third row) classes. These 3 sets of hand shapes correspond to the first three peaks of the performance index. These were obtained with 500 (red line) and 1000 (blue line) K-means iterations. Although the absolute maximum of the index suggests 4 clusters, we observed that the solution corresponding to 7 clusters provided us with three additional functional movements. If we consider the next index peak at 10 clusters, this solution gives additional states which are either a repetition of existing ones (S8 - previous S4 in the 7-classes solution) or noise (non-functional hand shapes such as S9, S10). For this reason, in what follows, we selected the solution corresponding to 7 clusters.

**Figure S4.**
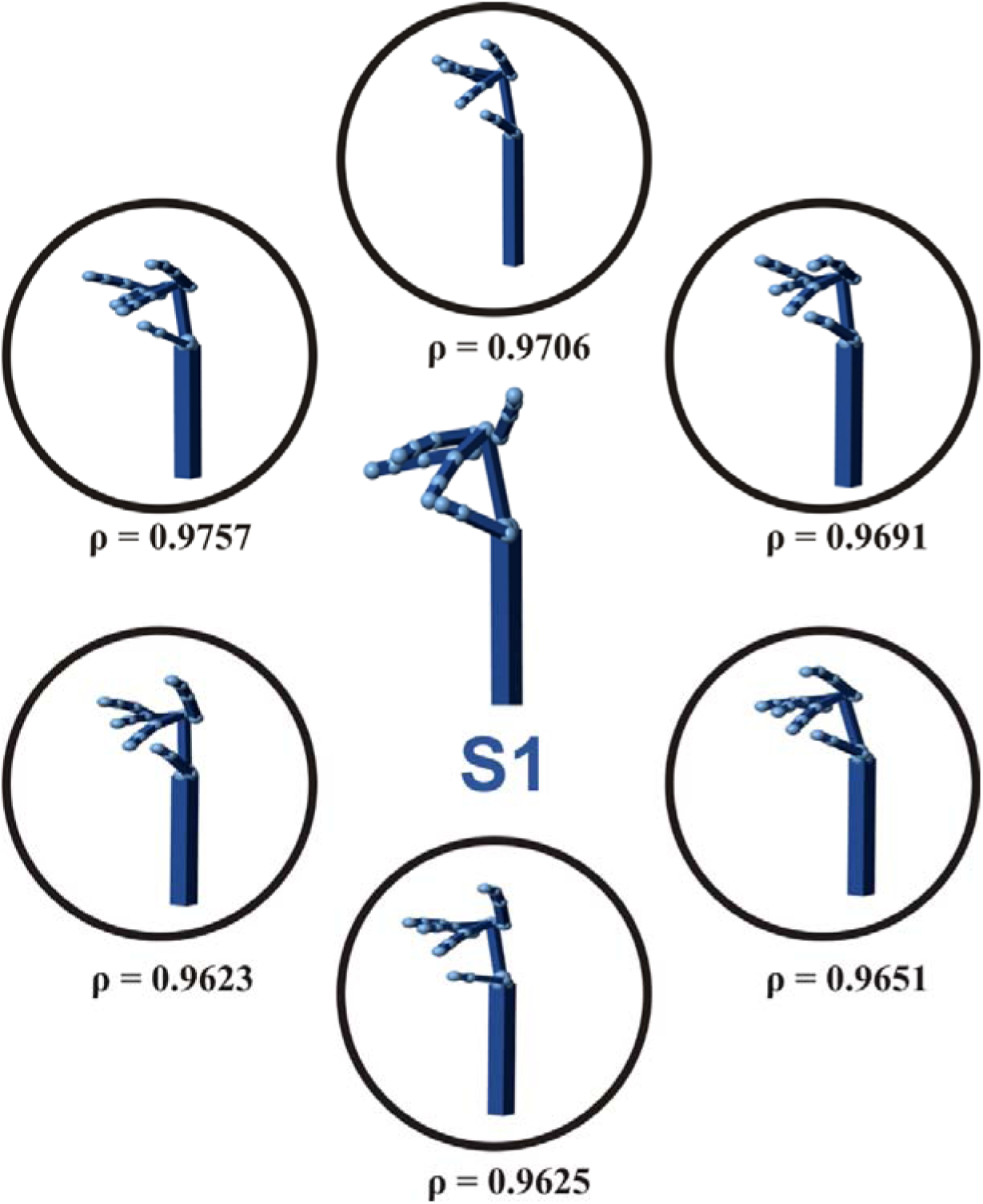
Variability of hand configurations within cluster S1. To interpret more accurately S1, we show the variability of the hand configurations within its cluster. Specifically, we show the first six configurations ranked by their correlation (*ρ*) with the centroid. It can be noted a certain excursion from the average for the index and little finger positions, while the motor configuration of the middle finger is consistent.

**Table S1.** **Everyday objects used by participants.** The diameter (or width for non-cylindrical objects), height and weight are reported. Items are sorted according to increasing diameter.

| OBJECT | DIAMETER<br>OR<br>WIDTH (cm) | HEIGHT (cm) | WEIGHT (g) |
| --- | --- | --- | --- |
| toothpick | 0.2 | 6.5 | 0.1 |
| coffee cup (handle) | 0.9 | 3.5 | 73 |
| teaspoon | 1 | 12 | 2 |
| knife | 1 | 16 | 3 |
| spoon | 1.2 | 17 | 3 |
| fork | 1.2 | 18 | 5 |
| cup (handle) | 1.7 | 6.5 | 426 |
| egg | 2.2 | 7 | 44 |
| jug (handle) | 2.9 | 13 | 95 |
| frying pan (handle) | 3 | 16 | 270 |
| salt container | 4 | 9 | 100 |
| sachet of sugar | 4.4 | 7 | 6 |
| tea bag | 4.5 | 5.5 | 2 |
| glass | 7 | 8.5 | 2 |
| bottle of water | 7 | 35 | 20 |
| packet of cigarettes | 7 | 5 | 4 |
| circular ashtray | 7.4 | 4.3 | 72 |
| jar | 8 | 7.5 | 193 |
| napkin | 19.5 | 19.5 | 4 |
| plate | 22 | 1.5 | 12 |
| placemat | 43 | 30 | 126 |

**Table S2.**
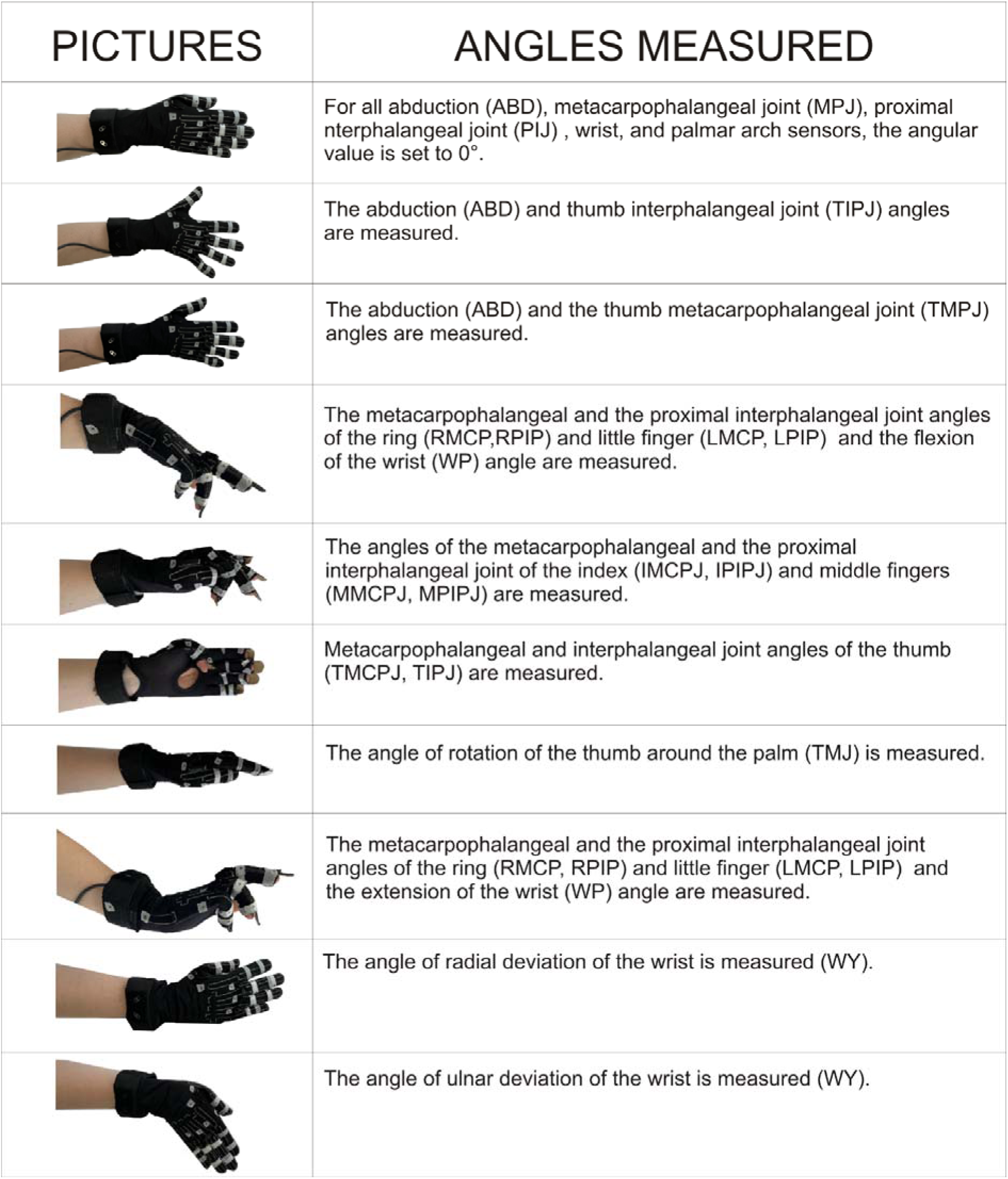
**Angles measured for each reference hand posture.**

| PICTURES | ANGLES MEASURED |
| --- | --- |
|  | For all abduction (ABD), metacarpophalangeal joint (MPJ), proximal interphalangeal joint (PIJ) , wrist, and palmar arch sensors, the angular value is set to 0°. |
|  | The abduction (ABD) and thumb interphalangeal joint (TIPJ) angles are measured. |
|  | The abduction (ABD) and the thumb metacarpophalangeal joint (TMPJ) angles are measured. |
|  | The metacarpophalangeal and the proximal interphalangeal joint angles of the ring (RMCP,RPIP) and little finger (LMCP, LPIP) and the flexion of the wrist (WP) angle are measured. |
|  | The angles of the metacarpophalangeal and the proximal interphalangeal joint of the index (IMCPJ, IPIPJ) and middle fingers (MMCPJ, MPIPJ) are measured. |
|  | Metacarpophalangeal and interphalangeal joint angles of the thumb (TMCPJ, TIPJ) are measured. |
|  | The angle of rotation of the thumb around the palm (TMJ) is measured. |
|  | The metacarpophalangeal and the proximal interphalangeal joint angles of the ring (RMCP, RPIP) and little finger (LMCP, LPIP) and the extension of the wrist (WP) angle are measured. |
|  | The angle of radial deviation of the wrist is measured (WY). |
|  | The angle of ulnar deviation of the wrist is measured (WY). |

